# Chromosome-Specific Expansion and Diversification of the Thionin Gene Family in Barley

**DOI:** 10.64898/2026.06.19.733385

**Authors:** Yao Fu, Joanne Russell, Miriam Schreiber, Jorunn I. B. Bos

## Abstract

Thionins are cysteine-rich peptides involved in plant defense. However, their genomic organization, evolutionary expansion, and potential function in barley remain unclear. Here, we integrated reference genome, pan-genome, and pan-transcriptome resources to systematically characterize the thionin gene family in barley. Fifty-six thionin genes were identified in the reference genome Morex V3, displaying pronounced chromosomal clustering and high sequence conservation consistent with extensive tandem duplication. Promoter analysis of these genes revealed enrichment of *cis*-acting elements associated with stress- and hormone-related signaling pathways, suggesting a potential role for thionins in biotic stress responses, including aphid defense, as suggested by previous studies. Analysis of 20 barley genotypes revealed substantial copy number variation, particularly on chromosomes 6H and 7H, indicating dynamic population-level expansion. Sequence-based clustering grouped thionins into ten clusters and five singletons, with major clusters corresponding to specific chromosomes. Integration of pan-transcriptome data showed that transcriptional activity was largely confined to four major clusters. Aphid infestation of four genotypes featuring copy number variation in chr 6H thionin genes resulted in a strong induction of thionin gene expression, with more pronounced responses during poor-host interactions. Aphid-induced expression tended to increase with thionin gene number, however, this no correlation was observed regarding basal gene expression levels. Together, these findings indicate that the thionin gene family in barley has undergone species-specific expansion driven by tandem duplication and contributes to genotype-dependent aphid defense responses.

## INTRODUCTION

Thionins are small cysteine-rich peptide proteins found exclusively in plants. They typically contain six to eight cysteine residues and are characterized by two consecutive cysteines at positions three and four (Bohlmann & Apel, 1991; Bohlmann & Boekaert, 1994; Iwai et al., 2002; Muramoto et al., 2012). The first thionin protein, purothionin, was isolated from wheat endosperm in 1942 and represents one of the earliest identified plant antimicrobial peptides (Balls & Hale, 1942). Since then, thionins have been identified in a wide range of plant species. Based on structural features, thionins can be classified into α/β-thionins and γ-thionins. As γ-thionins share strong structural similarity with defensins, they are now more commonly referred to as plant defensins. Despite structural differences, both groups display antimicrobial and antifungal activities in vitro. α/β-thionins are widely distributed across plant species but have been most extensively studied in grasses, particularly in wheat and barley. In *Arabidopsis thaliana*, four thionin genes have been identified, among which Thi2.1 confers resistance to *Fusarium oxysporum f. sp. matthiolae* (Epple et al., 1998). Transgenic expression of Thi2.1 in tomato enhances resistance to several pathogens (Chan et al., 2005), while recombinant thionin peptides expressed in mammalian endothelial cell systems inhibit the growth of bacteria, including *Escherichia coli* and *Staphylococcus aureu* (Loeza-Angeles et al., 2008). In rice, 44 thionin genes have been identified. Functional studies have shown that overexpression of OsThi9 reduces cadmium accumulation and toxicity, while OsTHI7 decreases susceptibility to the pathogens *Meloidogyne graminicola*and *Pythium graminicola*. Recombinant OsTHION15 also exhibits inhibitory activity against several pathogens (Ji et al., 2015; Boonpa et al., 2019; Liu et al., 2023). In wheat, thionins are mainly represented by purothionins that accumulate predominantly in the endosperm of developing seeds. Injection of purified purothionin into the hemolymph of *Manduca sexta* larvae results in approximately 50% mortality (Kramer et al., 1979). Expression of β-purothionin in Arabidopsis suppresses fungal growth and stomatal infection on leaves (Oard et al., 2006), and exogenous application of β-purothionin or purothionin-α2 reduces the severity of leaf rust infection caused by *Puccinia triticina* in bread wheat (*Triticum aestivum* L.) (Panthi et al., 2024).

In barley, thionins comprise two main groups: a small gene family encoding endosperm-specific hordothionins, and a large multigene family encoding leaf thionins (Höng et al., 2021). Mature leaf thionins accumulate in cell walls and vacuoles (Bohlmann et al., 1988; Reimann-Philipp et al., 1989), exhibit photoreactivity (Reimann-Philipp et al., 1989), and can be induced by various elicitors (Fischer et al., 1989; Andresen et al., 1992). Similar to thionins in other plant species, barley leaf thionins exhibit strong antimicrobial and antifungal activities in vitro (Bohlmann et al., 1988; Höng et al., 2021). However, direct evidence for their functional roles in plant defense in vivo remains limited. Recent studies suggested that thionins may contribute to barley resistance against aphids (Escudero-Martinez et al., 2017). Transcriptomic analyses identified several thionin genes that were strongly induced during aphid infestation (Delp et al., 2009). Some of these genes showed significant upregulation during poor-host interactions with *Myzus persicae* (*M.persicae*) and, to a lesser extent, during host interactions with *Rhopalosiphum padi* (*R.padi*). Furthermore, ectopic expression of two barley thionin genes in *Nicotiana benthamiana* (*N.benthamiana*) reduced host susceptibility to *M. persicae* (Escudero-Martinez et al., 2017). In the partially aphid-resistant wild barley accession *Hordeum spontaneum* 5 (HSP5), expression of three thionin genes increased more than sevenfold within 24 hours following aphid infestation (Leybourne et al., 2019).

Despite these findings, detailed functional analysis of the barley thionin gene family remains challenging. The large number of thionin genes and their high sequence similarity complicate the discrimination and functional characterization of individual family members, for example through gene editing approaches (Holtorf et al., 1995; Escudero-Martinez et al., 2017). Earlier studies also suggested substantial variation in thionin gene copy number among barley cultivars, as indicated by differences in southern blot hybridization signals (Bunge et al., 1992). However, the lack of high quality long-read genome-scale sequencing data at that time prevented a more detailed investigation of this variation. As a result, the extent of sequence diversity among thionin genes, the variation in gene copy numbers among barley genotypes, and the functional consequences of this variation remain largely unclear. The recent development of barley pan-genomic and pan-transcriptomic resources now provides powerful tools to address these questions. Analysis of the barley pan-genome revealed extensive copy number variation (CNV) at the thionin locus across different genotypes (Jayakodi et al., 2024). Among 76 barley genotypes analyzed, the total number of thionin genes exceeded 500, with individual genotypes containing between three and seventy-six copies. In addition, a barley pan-transcriptome comprising 20 inbred genotypes has recently been established, providing expression profiles for genes across multiple tissues and developmental stages (Guo et al., 2025). These resources offer new opportunities to systematically investigate the diversity, evolution, and expression patterns of the thionin gene family in barley.

In this study, we utilized the barley reference genome Morex V3 together with pan-genomic and pan-transcriptomic data from 20 barley genotypes (Mascher et al., 2021; Jayakodi et al., 2024; Guo et al., 2025) to systematically catalogue and characterise the genomic organization of the thionin gene family. We characterized the genomic distribution, CNV, and evolutionary relationships of thionin genes across 20 barley genotypes and investigated their expression patterns across tissues. We further assessed thionin expression responses to aphid infestation in four genotypes exhibiting pronounced CNV differences. These analyses provide new insights into the structural diversity, evolutionary dynamics, and potential functional roles of thionins in barley defense responses.

## MATERIALS AND METHODS

### Identification of thionins in Morex V3

To identify members of the thionin gene family in barley, protein sequences derived from the Morex V3 reference genome (Mascher et al., 2021) were retrieved and used as the local protein database. A comprehensive thionin reference dataset was constructed by integrating 4 experimentally characterized *Arabidopsis* thionins (Sels et al., 2008; Almaghrabi et al., 2019), 39 rice thionins (Boonpa et al., 2019), and 133 recently reported thionin sequences mined from transcriptomes of more than 1,000 plant species (Höng et al., 2021). BLASTP searches were performed against the barley proteome using the combined thionin dataset as query sequences, with an E-value threshold of 1×10^−5^. In parallel, a hidden Markov model (HMM) profile corresponding to plant thionins was retrieved from the Pfam database (http://pfam-legacy.xfam.org/) and used as the query for HMMER v3.0 (Eddy et al., 2021) searches against the Morex V3 protein dataset, applying the same *E*-value threshold.

Candidate thionin proteins identified by both BLASTP and HMMER were retained for further analysis. Redundant protein isoforms originating from alternative splicing of the same gene locus were removed, resulting in a non-redundant candidate set. Multiple sequence alignment of the retained thionin proteins was conducted using MAFFT v7 (Katoh et al., 2013). Conserved thionin domains were identified based on the alignment and previously reported domain features (Azmi et al., 2021). Poorly aligned or anomalous regions were manually trimmed to improve alignment quality and downstream analytical accuracy.

### Protein physicochemical property prediction

Theoretical molecular weight (MW), isoelectric point (pI), and other physicochemical parameters of the predicted thionin proteins were calculated using the ProtParam tool available on the ExPASy server (https://web.expasy.org/protparam/).

### Signal peptide, conserved domain, and motif analysis

Signal peptide sequences were predicted using SignalP v6.0 (Nielsen et al., 2024). Conserved domains were annotated using the NCBI Conserved Domain Database (CDD) (Marchler-Bauer et al., 2025). Conserved motifs were identified using MEME Suite v5.5.9 (Bailey et al., 2015) with default parameters, and only motifs with E-value lower than 1×10^−5^ were retained for subsequent analysis.

### Multiple sequence alignment and phylogenetic tree construction

Thionin protein sequences were aligned using MAFFT v7, followed by manual removal of gap-rich and poorly aligned regions. The best-fit amino acid substitution model was selected using ModelFinder implemented in IQ-TREE v3.0.1 (Wong et al., 2026) based on the Bayesian Information Criterion (BIC), and the JTT+G4 model was identified as the optimal model. Maximum-likelihood (ML) phylogenetic trees were then reconstructed in IQ-TREE using the selected model with 1000 standard bootstrap replicates to assess branch support. The resulting phylogenetic tree was visualized and annotated using iTOL (Letunic et al., 2024), with bootstrap support values ≥ 50% shown at the corresponding nodes.

### Chromosomal distribution, gene structure, and synteny analysis

Genomic coordinates of the identified thionin genes were extracted from the corresponding GFF annotation files. Chromosomal localization, gene structure visualization, and synteny analysis were conducted using TBtools-II v1.106 (Chen et al., 2023), with syntenic relationships identified using the integrated One-Step MCScanX module.

### Sequence identity analysis

To assess sequence similarity among thionin family members, nucleotide and protein sequences were aligned using MAFFT v7 with default parameters. Pairwise sequence identity matrices were generated, and results were visualized using TBtools-II.

### *Cis*-acting regulatory element analysis

The upstream regulatory regions of the candidate thionin genes were extracted using TBtools-II. For each gene, a 2-kb genomic sequence upstream of the translation start codon (ATG) was retrieved from the barley Morex V3 reference genome. The extracted promoter sequences were subsequently analyzed using the PlantCARE database (Lescot et al., 2002) to identify putative *cis*-acting regulatory elements (*cis*-elements). All analyses were performed with the default parameters provided by PlantCARE. Visualization of the predicted *cis*-elements was carried out using Python v3.11.13.

### Identification and curation of thionin genes across 20 barley genotypes

Twenty genomes were selected from the 76 barley genotypes included in the barley pan-genome (Jayakodi et al., 2024) based on their availability in the barley pan-transcriptome dataset (Guo et al., 2025), thereby enabling downstream extraction of gene expression information for the identified thionin genes. Thionin candidates were identified across the 20 genomes using a combined BLASTP and HMMER-based approach. Protein sequences of thionins identified from the barley reference genome Morex V3 were used as queries in BLASTP searches with the following criteria: *E*-value < 1×10^-5^, mismatches ≤ 10, and alignment coverage > 70%. In parallel, HMMER v3.0 was employed to search each genome using a thionin HMM, with an *E*-value cutoff of < 1×10^-5^. Only protein sequences detected by both BLASTP and HMMER were retained as high-confidence thionin candidates. All identified candidates were subsequently compiled and grouped according to their chromosomal locations.

To assess sequence integrity, candidate protein sequences were aligned using MAFFT v7 and manually curated by comparison with the characteristic thionin structural features observed on different chromosomes in the Morex V3 reference genome. Misaligned or irregular regions were trimmed to restore typical thionin architecture. For candidates located on unanchored scaffolds (chrUn), chromosomal origin was inferred based on sequence similarity and structural characteristics. The curated protein sequences were subsequently used as templates to refine their corresponding coding sequences (CDSs), ensuring consistency between CDSs and protein structures.

### Phylogenetic analysis across 20 barley genotypes

The trimmed CDSs of candidate thionin genes identified across the 20 barley genotypes were subjected to ML model testing in IQ-TREE v3.0.1. The best-fit nucleotide substitution model was selected using ModelFinder based on the BIC, with TVM+R3 identified as the optimal model. ML phylogenetic trees were subsequently reconstructed under the selected model, and branch support was evaluated using 1000 ultrafast bootstrap replicates (UFBoot). The resulting phylogenetic tree was visualized using iTOL, and only branches with bootstrap support values ≥ 70% were displayed.

### Clustering and CNV analysis based on CDS identity

CDSs of candidate thionin genes from the 20 barley genotypes were aligned using MAFFT v7 with the codon alignment option (--codon) to preserve reading frame integrity. A pairwise nucleotide identity matrix was generated from the resulting alignment, and sequences sharing ≥ 90% identity were assigned to the same cluster. Sequences that did not meet this threshold with any other sequence were defined as singletons. For each genotype, the number of sequences assigned to each cluster was quantified to assess CNV. Within individual genotypes, sequences that were 100% identical at the CDS level were further grouped into subclusters, and the number of subclusters per genotype was recorded.

For downstream visualization of divergence among clusters, pairwise *p*-distances were calculated from the identity matrix (Equation 1). For two clusters *C_i_* and *C_j_*, the inter-cluster distance was defined as the mean p-distance across all pairwise comparisons between sequences belonging to the two clusters (Equation 2).

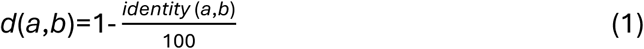

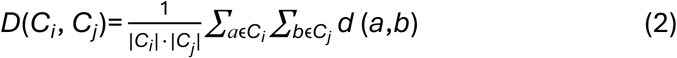

For comparisons between a cluster and a singleton, the distance was calculated as the mean *p*-distance between the singleton sequence and all members of the cluster. For two singletons, distances were obtained directly from the pairwise *p*-distance matrix. Cluster relationship networks and sequence identity heatmaps were generated using Python v3.11.13.

### Identification and curation of thionin transcripts across 20 barley genotypes

Transcriptome assemblies and corresponding expression datasets for 20 barley genotypes were retrieved from the barley pan-transcriptomic resource (Guo et al., 2025). To identify thionin genes represented in these transcriptomes, curated thionin CDSs derived from the barley pan-genome were used as query sequences. For each genotype, a BLASTN database was constructed from its pan-transcriptome assembly, and sequence similarity searches were performed using the following criteria: *E*-value ≤ 1×10^-5^, alignment length ≥ 200 bp, and mismatches ≤ 10. BLAST hits were subsequently processed by normalizing transcript identifiers (i.e., removal of isoform suffixes) and removing redundant entries, yielding a preliminary set of thionin candidates for each genotype. To ensure accurate genomic assignment, positional information for the candidate transcripts was obtained from genotype-specific pan-transcriptome GTF files, and corresponding genomic coordinates were retrieved from the pan-genome GFF annotation. Only candidates exhibiting concordant chromosomal identity, consistent strand orientation, and overlapping genomic intervals between transcript-based and genome-based annotations were retained. All isoforms associated with the retained genes were collected and grouped by chromosome for subsequent sequence curation. Refinement of thionin transcript sequences was guided by conserved thionin CDS characteristics established from Morex V3. Candidate sequences were aligned using MAFFT v7, and manual trimming was performed to remove intronic remnants, incomplete exons, and any aberrant sequence extensions. Redundant isoforms were removed, and distinct isoforms were preserved. If multiple plausible isoforms existed for a gene, all were retained for downstream analyses. To further validate their coding potential, all curated sequences underwent systematic open reading frame (ORF) detection and translation. ORFs were scanned in all three reading frames using ATG as the start codon and TAA, TAG, or TGA as stop codons. Regions containing a valid start-stop codon pair were designated as functional ORFs and translated into amino acid sequences using the standard codon table. Sequences lacking a biologically valid ORF were classified as putative pseudogenes. For these, full-length triplet translation was performed to recover potential coding information, and their pseudogene status was documented.

### Cluster assignment and subcluster definition of thionin transcripts

To assign barley pan-transcriptome transcripts to the thionin clusters previously defined in the pan-genome, BLASTN searches were performed against the curated thionin CDSs derived from the pan-genome. Because a subset of transcripts was incomplete, candidate sequences were first divided into two categories based on transcript length: standard-length candidates (≥ 300 bp) and incomplete or short candidates (< 300 bp). For standard-length candidates, BLAST hits were retained only when all of the following criteria were satisfied: nucleotide sequence identity (pident) ≥ 90%, transcript query coverage (qcov) ≥ 70%, and CDS subject coverage (scov) ≥ 70%. For incomplete or short candidates, which may represent authentic but truncated transcript isoforms, more stringent thresholds were applied for sequence identity and query coverage (pident ≥ 90% and qcov ≥ 90%). In contrast, no minimum CDS coverage threshold was imposed for these candidates in order to avoid excluding valid partial homologs. For each transcript, only the BLAST hit with the highest bitscore that satisfied the above criteria was retained as the best-matching CDS. The cluster identity of the matched CDS was then transferred to the corresponding transcript, thereby assigning each transcript to a predefined thionin cluster. The number of transcripts assigned to each cluster was subsequently quantified across all 20 barley genotypes. All pan-transcriptome thionin sequences were then aligned using MAFFT v7 to generate a pairwise sequence identity matrix. Within each genotype, transcripts exhibiting 100% nucleotide sequence identity were defined as subclusters. The number of subclusters within each thionin cluster was calculated to assess intra-genotype redundancy and genotype-specific expansion patterns.

### Expression quantification across genotypes and tissues

Transcript expression levels were retrieved based on the candidate thionin transcript identifiers obtained from the pan-transcriptome. The pan-transcriptome dataset provides expression profiles for caryopsis, coleoptile, inflorescence, root, and shoot, with each tissue represented by three biological replicates (Guo et al., 2025). For genotypes HOR10350, HOR7552, and HOR8148, only two biological replicates were available for root tissue. For each tissue and genotype, the mean expression value across all available biological replicates was used for downstream analyses. As transcripts within the same subcluster share 100% sequence identity, expression values could not be unambiguously assigned to individual subcluster members. Therefore, expression values of all transcripts belonging to the same subcluster were summed to represent the total expression level of that subcluster, and the number of transcripts within each subcluster was recorded. In addition, total expression values were calculated for each thionin cluster within each genotype.

### Plant and aphid materials

Following genotype-level analyses, plant-level experiments were conducted using individual barley plants to investigate aphid-barley interactions. For this purpose, seeds of four barley genotypes: Akashinriki, HOR10350, HOR21599, and Morex were used for aphid infestation assays. Seeds were surface-sterilized prior to germination. Briefly, seeds were rinsed with distilled water, sterilized sequentially with 75% (v/v) ethanol and 2% (w/v) sodium hypochlorite, and then washed thoroughly with sterile distilled water. Sterilized seeds were placed on moistened filter paper and germinated at room temperature in darkness. After germination, seedlings were transplanted into pots containing bulrush compost mix (Bulrush, Northern Ireland) and grown in a controlled-environment growth room under an 16 h light / 8 h dark photoperiod, with a day/night temperature regime of 22 / 20 °C and 60% relative humidity. Seedlings were grown for 10 days until the first true leaf stage (Zadoks et al., 1974). Plants at the same developmental stage were selected for subsequent aphid infestation experiments.

Aphids were maintained in a growth chamber at 18 °C under a 16 h light / 8 h dark photoperiod and 50% relative humidity. Aphids were reared in ventilated plastic cups containing excised host plant leaves, with leaf bases immersed in water to maintain turgor. *R.padi* (JHI_JB; Thorpe et al., 2018) were maintained on leaves of barley (*Hordeum vulgare cv*. Optic), whereas *M.persicae* (JHI_genotype O; Thorpe et al., 2018) were maintained on leaves of *Brassica napus*.

### Aphid infestation of barley plants

Ten-day-old barley plants of the genotypes Akashinriki, HOR10350, HOR21599, and Morex with uniform growth were used for aphid infestation experiments. Three independent biological replicates were performed. Within each biological replicate, four treatment groups were established for each genotype: an untreated control, a clip-cage control in which clip cages were placed on leaves without aphid infestation, and two aphid infestation treatments involving *R.padi* or *M.persicae*. Each treatment group consisted of four individual plants per genotype. For each aphid infestation treatment, 20 mixed-instar aphids were randomly selected and placed on the leaf surface of each plant. Clip cages were used to confine aphids to the infestation site. All infestations were initiated at 10:00 a.m., and aphids were removed after 24 h. Following infestation, a leaf segment approximately 2 cm in length was excised from the aphid feeding site of each plant. Leaf segments of the same length and corresponding leaf position were also collected from clip-cage control and untreated control plants. Samples from the same treatment group, genotype, and biological replicate were pooled into a single 2 mL microcentrifuge tube, immediately frozen in liquid nitrogen, and stored at -80 °C until further analysis.

### RNA extraction, quantitative RT-PCR

Leaf samples were transferred to 2 mL microcentrifuge tubes containing 5 mm stainless steel beads and homogenized using a TissueLyser II (Qiagen, UK) at 25 Hz for 30 s, repeated for three cycles. Total RNA was extracted using TRIzol reagent (Invitrogen, UK) according to the manufacturer’s instructions, followed by removal of genomic DNA contamination using the TURBO DNA-free™ Kit (Invitrogen, UK). RNA concentration and purity were assessed using a NanoDrop ND-1000 spectrophotometer (Thermo Fisher Scientific, UK). First-strand cDNA was synthesized from DNase-treated RNA using the SuperScript III First-Strand Synthesis System (Sigma-Aldrich, UK) following the manufacturer’s protocol. RT-qPCR reactions were performed using PowerUp SYBR Green Master Mix (Applied Biosystems, UK) on a QuantStudio 3 Real-Time PCR System (Applied Biosystems, UK). Each 10 μL reaction mixture contained 5 μL SYBR Green Master Mix, gene-specific forward and reverse primers (final concentration 400 nM each), and approximately 10 ng of cDNA template. The thermal cycling conditions were as follows: 50 °C for 2 min for UDG activation, 95 °C for 2 min for initial denaturation, followed by 40 cycles of 95 °C for 15 s and 60 °C for 1 min. Fluorescence signals were collected at the end of each annealing/extension step. A melt curve analysis was performed after amplification to verify the specificity of PCR products using the following program: 95 °C for 15 s, 60 °C for 1 min, and 95 °C for 15 s, with continuous fluorescence acquisition. Each reaction was performed with three technical replicates.

Due to the high sequence similarity among CDSs of thionin gene family members, specific amplification of individual thionin genes is technically challenging. Therefore, a family-level expression analysis strategy was adopted, and consensus primers were designed to collectively target thionin genes located on chromosome 6H across the barley genotypes examined. To ensure primer design accuracy, CDSs of thionin genes located on chr 6H in the genotypes Akashinriki, HOR10350, HOR21599, and Morex were first subjected to quality filtering to remove incomplete or truncated sequences. Within each genotype, fully conserved regions shared by all retained sequences were identified and used as candidate sites for forward and reverse primer design. In the HOR21599 and Morex genotypes, the number of thionin genes on chr 6H was relatively high and sequence divergence was therefore higher, such that no fully conserved region across all sequences was suitable for both forward and reverse primer binding. Therefore, for these two genotypes, the most highly conserved regions were selected for forward primer design, resulting in two genotype-specific forward primers differing by three or four nucleotides, which were paired with a single common reverse primer. Accordingly, each of the HOR21599 and Morex genotypes was represented by two forward primers and one reverse primer to ensure comprehensive detection of thionin family expression. Actin (*ACT*), ubiquitin-conjugating enzyme (*UBC*), two commonly used reference genes, were included as internal controls for normalization. Primer sequences and design parameters are provided in Table S15.

Cycle threshold (Ct) values were obtained using QuantStudio 3 software. Technical replicates were subjected to quality control, and outliers were excluded when the standard deviation of Ct values exceeded 0.5. The remaining Ct values were averaged for subsequent analyses. Relative expression levels were calculated using the 2^−ΔΔCt^ method (Livak et al., 2001), with the untreated control group of Akashinriki used as the global calibrator for calculation. Relative expression values were further transformed into log_2_ fold-change values for downstream visualization and comparison. Summary statistics, including mean, median, standard deviation, standard error, and replicate number, were calculated from biological replicates. All data processing and statistical summaries were performed in Python v3.11.13.

## RESULTS

### Identification of thionin family members in the barley reference genome

Using a combined BLASTP- and HMMER-based strategy, a total of 56 putative thionin family members were identified from the barley reference genome Morex V3 based on their predicted amino acid sequences (Table S1). Multiple sequence alignment revealed that two candidates (HORVU.MOREX.r3.7HG0645080.1 and HORVU.MOREX.r3.7HG0645130.1) contained additional N-terminal amino acid sequences located upstream of the predicted signal peptide, likely resulting from inaccurate annotation or incomplete trimming of the gene models. These redundant regions were manually removed prior to subsequent analyses. In addition, eight candidates exhibited varying degrees of sequence truncation, indicating that the predicted genes encode incomplete thionin proteins.

To further characterize the 56 identified thionins, physicochemical properties, signal peptide predictions, and conserved domain annotations were systematically analyzed (Table S2-S3). The predicted protein lengths ranged from 86 to 137 amino acids (aa). Interestingly, intact thionins displayed chromosome-associated length patterns: proteins encoded on chr 1H were 133 or 136 aa in length, those on chr 6H were consistently 137 aa, and those on chr 7H were 134 aa. The predicted molecular weights ranged from 10.04 to 14.86 kDa, and the theoretical isoelectric points varied from 5.55 to 8.84, indicating substantial diversity in the physicochemical properties of barley thionins. All identified proteins contained the characteristic thionin domain (PF00321) according to the NCBI CDD, confirming their classification as members of the classical thionin family rather than the γ-thionin (plant defensin) group. Therefore, truncated sequences were retained for downstream analyses. Among the 56 candidates, four proteins were annotated as non-specific matches within the thionin domain superfamily, indicating that although they possessed the conserved thionin domain, they could not be confidently assigned to a specific subfamily. Notably, two of these proteins lacked a predicted signal peptide, likely due to incomplete N-terminal sequences.

### Phylogenetic analysis of barley thionin proteins reveals chromosome-associated clustering

To investigate the evolutionary relationships of barley thionins, we constructed a phylogenetic tree based on the amino acid sequences of 56 thionin proteins and further examined their motif patterns, sequence features, and conserved domains (Figure 1). The phylogenetic analysis resolved these proteins into four major groups (Figure 1A), with clustering patterns that largely correspond to the chromosomal locations of their encoding genes. Thionin proteins encoded by genes located on the same chromosome generally clustered together, whereas those encoded by genes from different chromosomes formed distinct branches. Due to the high sequence similarity among thionins, the phylogenetic tree showed limited resolution. Therefore, only nodes with bootstrap support ≥ 50 were considered. Groups I and II both comprised thionin proteins encoded by genes on chr 1H but formed two separate clades, indicating pronounced divergence among thionins originating from this chromosome. Group III contained all thionins encoded by genes on chr 6H, together with two proteins encoded by unanchored genes. These two proteins clustered closely with the chr 6H group, suggesting that their corresponding genes likely originated from chr 6H but were not properly anchored in the current genome assembly. Group IV included all thionins encoded by genes on chr 7H and one protein with an incomplete 86 aa sequence encoded by a gene on chr 5H. The placement of this truncated protein within the chr 7H cluster is likely due to phylogenetic bias caused by sequence truncation. Overall, the phylogenetic grouping closely mirrors the chromosomal distribution of the corresponding genes, suggesting that the expansion and diversification of the barley thionin family have largely followed chromosome-specific evolutionary trajectories.

**FIGURE 1.**
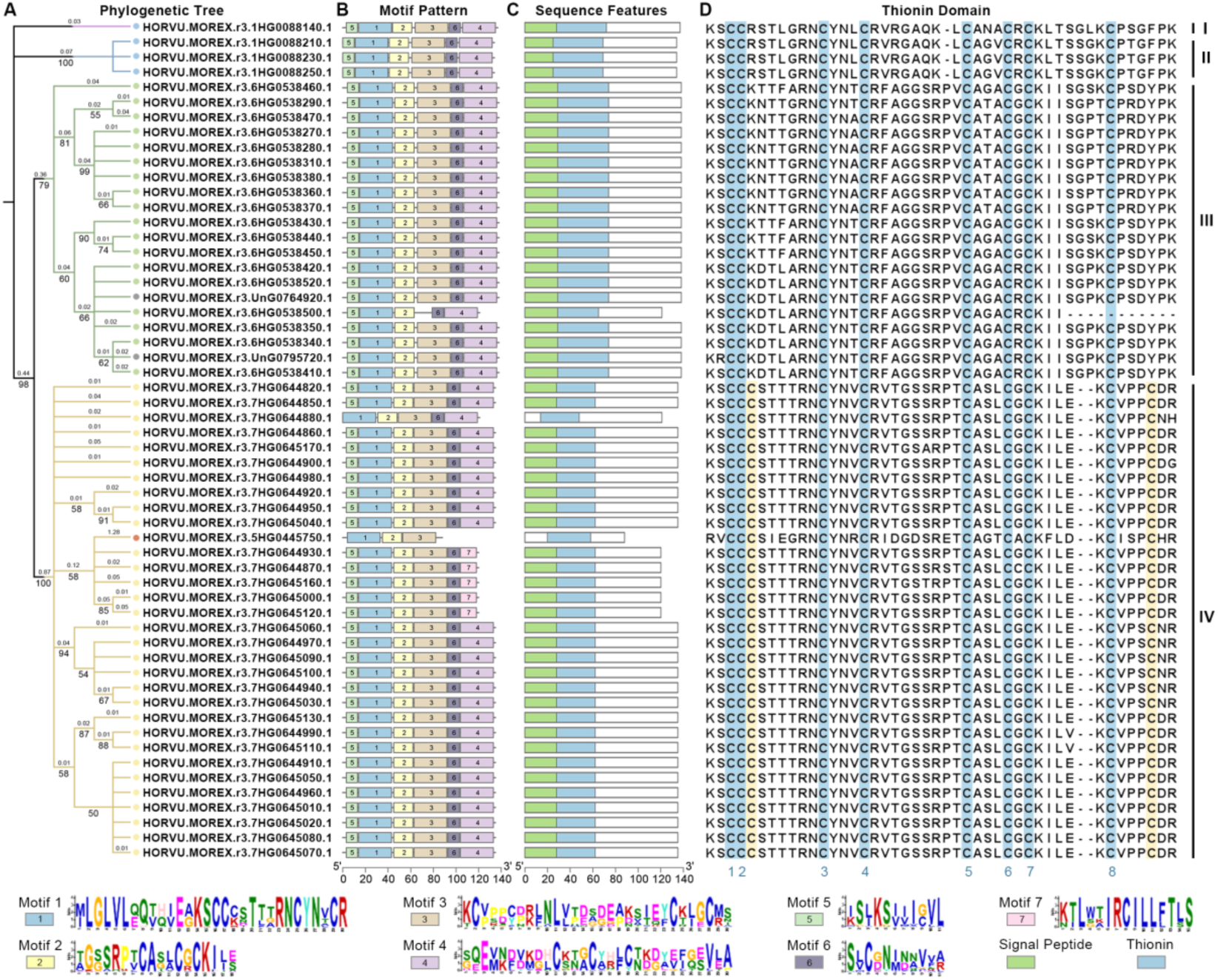
Phylogenetic relationships and architecture of conserved protein motifs of barley thionins. (A) Phylogenetic tree constructed from 56 barley thionin protein sequences. Branch lengths (numbers above the lines; only values > 0.01 are shown) and bootstrap support values (numbers below the lines; only values > 50 are shown) are indicated. Solid circles in corresponding colors denote the chromosomes on which each thionin gene is located. Blue, red, green, yellow, and gray represent chr 1H, 5H, 6H, 7H, and unknown, respectively. (B), (C) Motif composition and arrangement of thionins, shown as colored boxes. Protein lengths can be inferred from the scale at the bottom. Motif identities correspond to the legend. (D) Sequence analysis of the thionin domain. Cysteine residues are highlighted in blue and yellow, and the numbers in blue below the alignment indicate the total number of cysteine residues in all sequences.

Analysis of conserved domains and motif composition across the 56 identified thionin proteins revealed seven distinct motifs (Figure 1B). Among these, the motif arrangement “motif 5 - motif 1 - motif 2 - motif 3 - motif 6 - motif 4” was the most prevalent, occurring in 48 sequences. Overall, thionin proteins displayed highly conserved motif architectures, indicating a high degree of structural conservation across the family. Sequence feature analysis further demonstrated that all thionins contained the characteristic thionin domain (Figure 1C). Two members lacked a predicted signal peptide, most likely due to incomplete N-terminal sequences. Comparative analysis of the thionin domains showed group-specific variation in domain length (Figure 1D). Domains in Groups I and II comprised of 45 aa, those in Group III (excluding truncated sequences) comprised of 46 aa, whereas domains in Group IV comprised of 44 aa. In addition, cysteine residue composition varied among groups. Thionin domains in Groups I-III each contained eight cysteine residues, whereas Group IV domains harbored ten cysteine residues. Given that classical thionins typically contain six to eight cysteine residues, the presence of additional cysteines in Group IV suggests potential differences in disulfide bond configuration and domain architecture. Collectively, these observations suggest that Group IV thionins possess distinct domain features relative to the other groups.

The chromosomal distribution of thionin genes showed that members located on the same chromosome were typically arranged in tightly packed clusters within relatively restricted genomic regions (Figure 2A, Table S4). On chr 1H, four thionin genes were confined to a ∼187 kb region, with intergenic distances of approximately 50-83 kb, forming a small local cluster. A similar pattern was observed on chr 6H, where 18 thionin genes were distributed across a ∼1.3 Mb interval with intergenic distances ranging from 9 to 156 kb. The clustering on chr 7H was the most pronounced, with 33 thionin genes spanning a ∼0.9 Mb region and intergenic distances ranging from 5 to 198 kb. Overall, this dense local clustering, together with the predominance of short intergenic distances (mostly < 50 kb), suggests that the expansion of the barley thionin family was primarily driven by tandem and segmental duplication events. This is further supported by similarity analyses of the CDSs and amino acid sequences of the 56 thionins (Figure 2B, Table S5). The thionin sequences exhibited high sequence identity, especially among genes located on the same chromosome. For example, CDSs within chr 1H, 6H, and 7H shared over 85% identity. Amino acid sequences shared at least 75% identity, with most sequences sharing over 95% identity. Across all thionins, we identified 11 groups of genes with identical CDSs and 9 groups with identical amino acid sequences (Figure 2B, Table S5). Several genes with distinct CDSs encoded identical amino acid sequences, which explains why the number of identical protein sequence groups was smaller than the number of identical CDS groups. Taken together, the tight physical clustering of thionin genes, the highly conserved domain architectures, and the high sequence similarity among family members all point to extensive tandem and segmental duplication events that shaped the expansion of the thionin gene family in barley.

**FIGURE 2.**
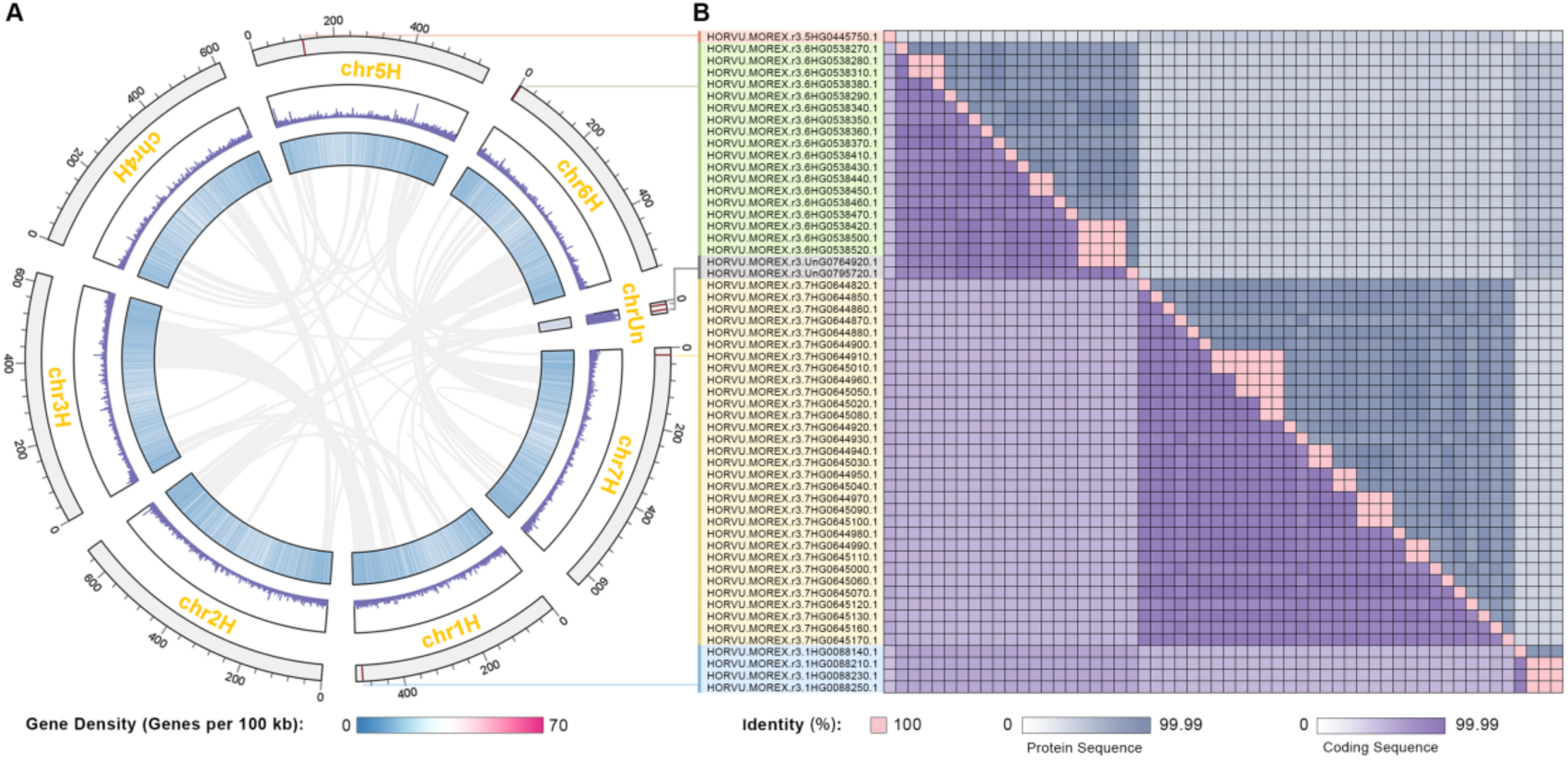
Chromosomal localization and sequence identity of barley thionin gene family. (A) Chromosomal map of the genomic positions of the 56 thionin genes. Genes are marked by the red line on the outermost circular track, with corresponding gene IDs labeled. Detailed genomic coordinates are provided in Table S4. The two inner circular tracks represent gene density on each chromosome, displayed as a heatmap and a line plot, respectively. Gray links in the center indicate syntenic relationships within the barley Morex V3 genome. (B) Sequence identity matrix of thionin CDS and protein sequences. Blue blocks represent protein sequence identity values from 0 to 99%, whereas purple blocks represent CDS sequence identity values from 0 to 99%. Pink blocks indicate sequences that are completely identical. Numerical values for sequence identity are provided in Table S5.

### Promoter *cis*-element analysis indicates potential roles of barley thionins in stress and hormone signaling

To examine the regulatory features of the identified thionin genes, *cis*-elements were analyzed within their promoter regions. Based on element abundance and functional annotation, 29 representative *cis*-elements were retained for subsequent analyses (Figure 3, Table S6, S7). Across all promoters, *cis*-elements were distributed relatively evenly, and overall composition patterns were highly similar among thionin genes located on different chromosomes (Figure 3A). The total number of *cis*-elements per gene ranged from 25 to 60. Among the analyzed genes, HORVU.MOREX.r3.1HG0088140.1 harbored the highest number of *cis*-elements, suggesting a potentially enhanced capacity for transcriptional regulation in response to environmental and hormonal signals. Functional categorization grouped the identified *cis*-elements into four major classes: stress response, hormone response, growth-related regulation, and light response (Figure 3B). Elements associated with stress and hormone responses were the most abundant across promoters, whereas light-responsive elements were also consistently detected.

**FIGURE 3.**
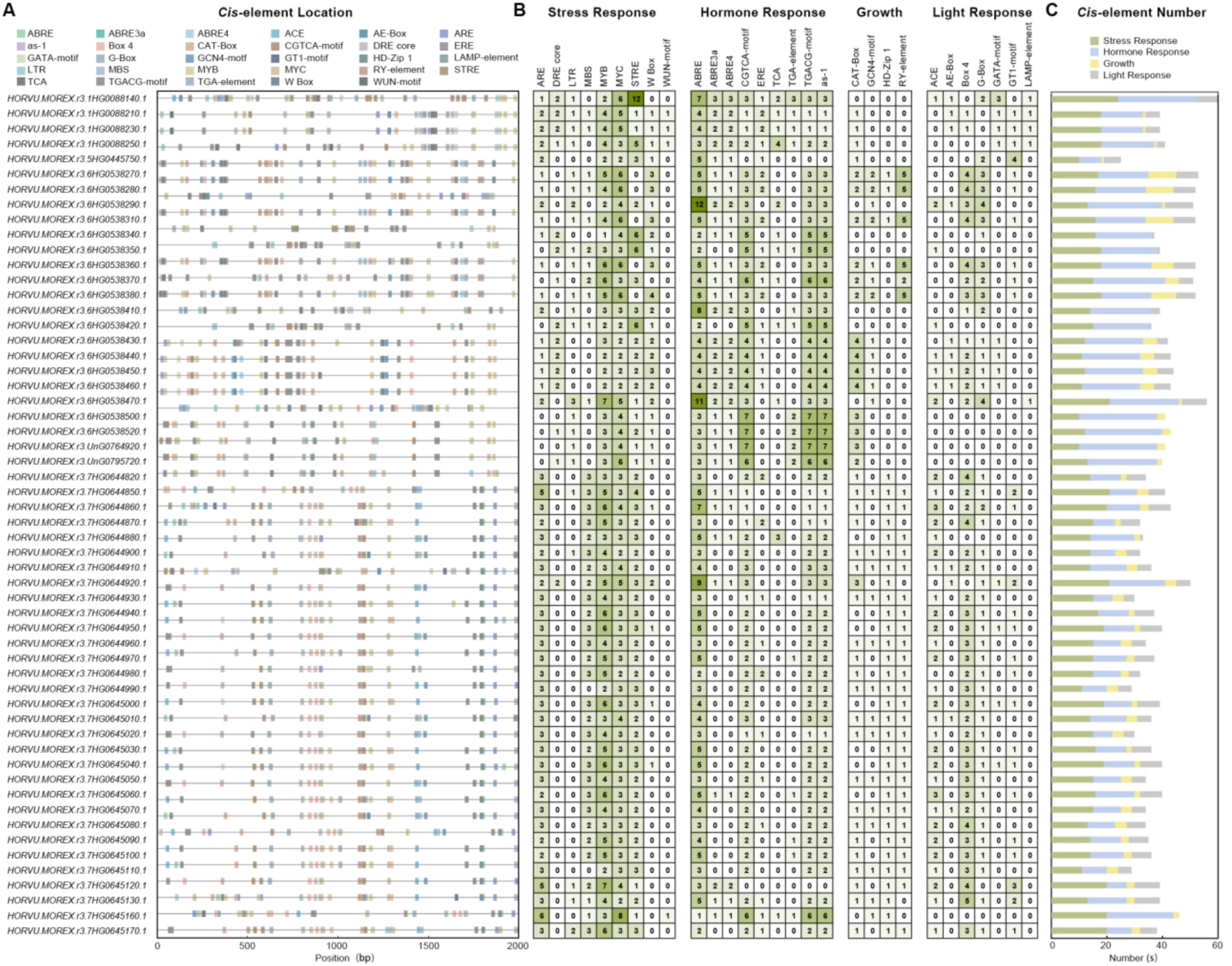
*Cis*-element location and number analysis of thionin genes. (A) Distribution of *cis*-elements across the promoter regions of 56 thionin genes. Different color blocks represent different *cis*-elements, and the horizontal axis indicates the length of the DNA sequence corresponding to the 2-kb upstream region of the CDS. (B) Analysis of *cis*-element numbers and categories in the promoter regions. Different colors and numbers indicate the abundance of various *cis*-elements in each gene promoter. (C) Total counts and relationships of *cis*-elements in each category. Green, blue, yellow, and gray blocks represent the categories of stress response, hormone response, growth, and light response, respectively.

Chromosome-associated differences in *cis*-element composition were observed (Figure 3C). Promoters of thionin genes located on chr 1H were enriched in stress-and hormone-responsive elements but contained few growth-related elements. In contrast, thionins on chr 6H showed a higher proportion of hormone-responsive elements, whereas thionins on chr 7H were relatively enriched in light-responsive elements. Within the stress-response category, MYB-binding site (MYB), MYC-binding site (MYC) and stress-responsive element (STRE) were the most frequently detected motifs. MYB motifs are recognized by MYB transcription factors and are widely involved in abiotic stress regulation. MYC motifs are associated with drought and abscisic acid (ABA) responses and commonly occur in stress-responsive promoters. STRE motifs are central to general stress signaling and were especially abundant in the promoter of HORVU.MOREX.r3.1HG0088140.1, which contained 12 such elements. Additional stress-related motifs included WRKY transcription factor binding sites (W-boxes) associated with wound and pathogen responses, Low-temperature-responsive elements (LTR) associated with cold stress and MYB binding site (MBS) and anaerobic response element (ARE) related to drought, oxidative stress and anaerobic conditions. Thionins on chr 7H showed noticeably higher numbers of ARE motifs than those on other chromosomes, suggesting potential chromosome-specific specialization. In the hormone-response category, all thionin promoters contained ABA-responsive elements (ABRE), which were also the most abundant hormone-related motifs. Promoters of thionins on chr 6H displayed higher numbers of activation sequence-1 (as-1), CGTCA-motif and TGACG-element, which are associated with salicylic acid (SA) responsiveness, oxidative stress and methyl jasmonate (MeJA) signaling. This pattern suggests that thionins on chr 6H may play important roles in plant defense pathways. Additional motifs in this category included ethylene-responsive elements (ERE), TGA-elements involved in auxin signaling and TCA-elements linked to SA responsiveness. Among growth-related elements, meristem expression elements (CAT-box) involved in meristem activity and RY elements associated with seed-specific regulation were the most common and occurred predominantly in thionins on chr 6H. For light-responsive elements, Box 4 and G-box were the most prevalent. Box 4 was widely distributed among thionins on chr 6H and chr 7H. G-box motifs respond to both light and jasmonic acid (JA) signaling and therefore may contribute to the involvement of thionins in defense and environmental adaptation.

In summary, the promoter regions of thionins in barley contain a large number of *cis*-elements related to plant defense and hormone responsiveness. This observation is in line with the role of thionins in barley defense mechanisms and resistance to stress (Bohlmann & Apel, 1991; Sels et al., 2008; Hao et al., 2016). Additionally, several *cis*-elements associated with light response suggest that the expression of thionin genes may be influenced or regulated by light conditions, in line with previous findings (Gausing *et al*., 1987; Reimann-Philipp et al., 1989a; Reimann-Philipp et al., 1989b).

### Pan-genome analysis reveals extensive CNV of thionin genes across 20 barley genotypes

Following the genome-wide characterization of thionin genes in the barley reference genome, which revealed extensive chromosomal clustering, tandem duplication, and a highly conserved sequence architecture, we further investigated the conservation and diversification of thionins across 20 barley genotypes in the pan-genome (Jayakodi et al., 2024). Using a combined BLASTP-and HMMER-based strategy followed by sequence curation and trimming, thionin family members were identified in each of the 20 genotypes (Figure 4, Table S8). The total number of thionin genes varied markedly among genotypes (Figure 4A). HOR3081 harbored only 22 thionins, whereas ZDM01467 contained as many as 83, indicating substantial genotype-specific expansion of the thionin family. Analysis of chromosomal distribution revealed contrasting patterns among chromosomes. Thionins located on chr 1H were relatively conserved, with two to four members present in all genotypes except HOR10350, in which no thionin genes were detected on chr 1H (Figure 4A). In contrast, thionins located on chr 6H and chr 7H exhibited much greater variability, ranging from three to 47 and from one to 52 members, respectively. These pronounced differences suggest that thionin genes on chr 6H and chr 7H have experienced frequent gene gain and loss events during barley genome evolution.

**FIGURE 4.**
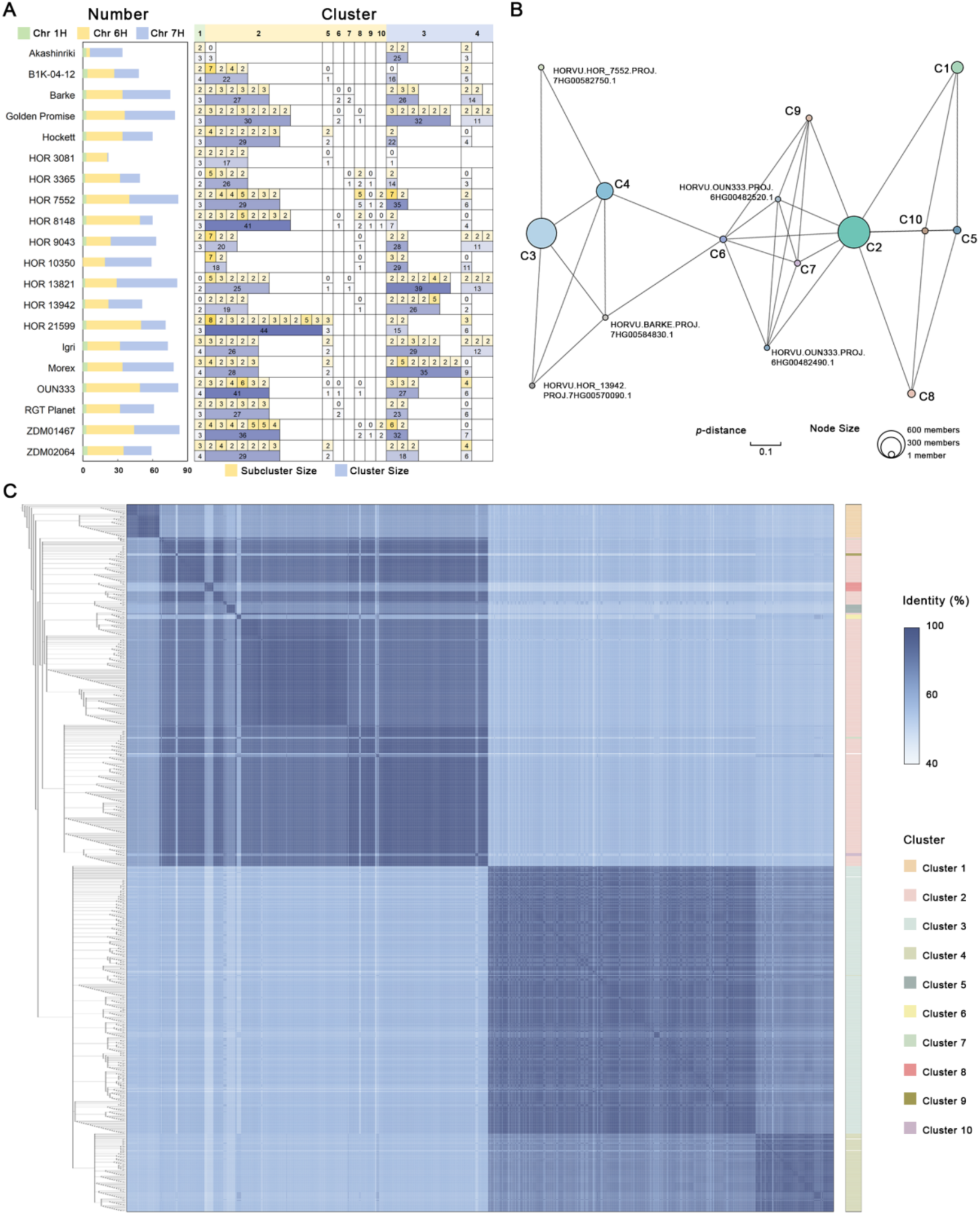
Numbers, clustering patterns, and sequence identity relationships of thionin genes across 20 barley genotypes. (A) Distribution of thionin numbers, cluster and subcluster composition across 20 barley genotypes. In the stacked bar plot, green, yellow, and blue indicate thionins located on chr 1H, chr 6H, and chr 7H, respectively. The heatmap on the right displays the number of members assigned to each cluster and subcluster in every genotype. Cluster sizes are represented by a blue color gradient, whereas subcluster sizes are shown using a yellow gradient. In the header, green, yellow, and blue shading indicate the chromosomal origins (chr 1H, chr 6H, chr 7H) of each cluster. (B) Network of sequence relationships among thionin clusters. C1-10 represent Clusters 1-10. Nodes with different colors represent the ten clusters and five singleton sequences, and node size reflects the number of members in each cluster. Edges indicate *p*-distances between representative sequences of each cluster. Detailed numerical values are provided in Table S11. (C) Phylogenetic tree and pairwise sequence identity heatmap of thionins from 20 barley genotypes. The heatmap displays sequence identity values using a blue color gradient, with detailed numeric values available in Table S9. The phylogenetic tree shows branches with bootstrap support ≥70. A color bar on the right indicates cluster assignments.

A phylogenetic tree constructed using all thionin CDSs from the 20 barley genotypes, together with pairwise sequence identity analyses (Figure 4C, Table S9), revealed that sequence variation among thionins was primarily associated with chromosomal origin rather than genotype. Thionins encoded on the same chromosome clustered together and exhibited high sequence similarity across genotypes. Using a 90% nucleotide sequence shared identity threshold, all thionin CDSs were grouped into 10 clusters and five singleton sequences (Figure 4C, Table S10). Clusters 1, 2, and 3 were the dominant groups, comprising 59, 537, and 479 members, respectively, and corresponded to thionins located on chr 1H, 6H, and 7H. Thionins on chr 1H formed a single cluster, indicating strong sequence conservation across genotypes. In contrast, thionins on chr 6H were distributed among seven clusters (Cluster 2, 5-10), whereas those on chr 7H were assigned to two clusters (Cluster 3, 4), reflecting greater sequence diversification in these chromosomes. Notably, thionins on chr 6H and 7H were predominantly assigned to Clusters 2 and 3, respectively. Pairwise *p*-distance analysis (Figure 4B, Table S11) showed that clusters originating from the same chromosome generally exhibited lower sequence divergence (approximately 0.13-0.17) than clusters from different chromosomes (approximately 0.28-0.37), indicating that sequence diversification among barley thionins is largely associated with chromosomal origin.

To evaluate expansion patterns, the number of members in each cluster was quantified for each genotype, and subclusters defined by 100% identical CDS sequences were further identified (Figure 4A, Table S10). On chr 1H, cluster 1 consisted mainly of one highly conserved thionin gene and one gene that underwent limited duplication, resulting in only a small number of subclusters. In contrast, cluster 2 on chr 6H displayed pronounced variation among genotypes, with member numbers ranging from 3 to 44. All genotypes except Akashinriki contained multiple subclusters within cluster 2, and HOR21599 exhibited the most extensive expansion, indicative of strong tandem duplication. Cluster 3 on chr 7H showed a similar pattern, with substantial variation in member numbers among genotypes, identifying chr 7H as another major hotspot of thionin family expansion.

Overall, clustering patterns reflect chromosomal-level sequence diversification, whereas variation within clusters reflects genotype-specific duplication. The limited clustering on chr 1H indicates strong sequence conservation, whereas the more complex clustering and subclustering patterns on chr 6H and 7H suggest increased sequence diversification and duplication, potentially contributing to functional divergence.

### Pan-transcriptome analysis reveals genotype-dependent and chromosome-specific expression patterns of barley thionins

The extensive copy numbers and CNV of the thionin gene family in the barley genome suggests that this gene family may play diverse roles across tissues and developmental stages. To systematically characterize thionin expression patterns, we analyzed their transcriptional profiles across multiple tissues and genotypes using the barley pan-transcriptome dataset reported by Guo et al (2025). Gene annotation systems differ between the pan-genome and pan-transcriptome datasets. We therefore combined BLASTN-based sequence similarity searches with chromosomal position information to identify pan-transcriptomic transcripts corresponding to pan-genomic thionin genes. Identified transcripts were curated, trimmed, and translated into corresponding amino acid sequences (Table S12). As shown in Figure 5A, the number of thionin transcripts detected in the pan-transcriptome varied markedly among genotypes, ranging from 6 to 48. Comparison of thionin copy numbers between the pan-genome and pan-transcriptome revealed clear chromosome-associated differences in transcript detection. On chr 1H, thionin numbers were nearly identical between the two datasets, indicating that thionins on this chromosome are broadly expressed under normal growth conditions. In contrast, thionins located on chr 7H were detected in the transcriptomes of only four genotypes, suggesting that their expression is genotype dependent or condition specific. On chr 6H, the number of thionins detected in the pan-transcriptome was slightly lower than in the pan-genome but still encompassed most annotated members, indicating generally high expression potential. In several genotypes, including Golden Promise, HOR13942, Igri, and RGT Planet, the number of thionins detected in the pan-transcriptome exceeded that in the pan-genome for specific chromosomes. This discrepancy is likely attributable to differences in annotation strategies that resulted in gene splitting and was not further examined here.

**FIGURE 5.**
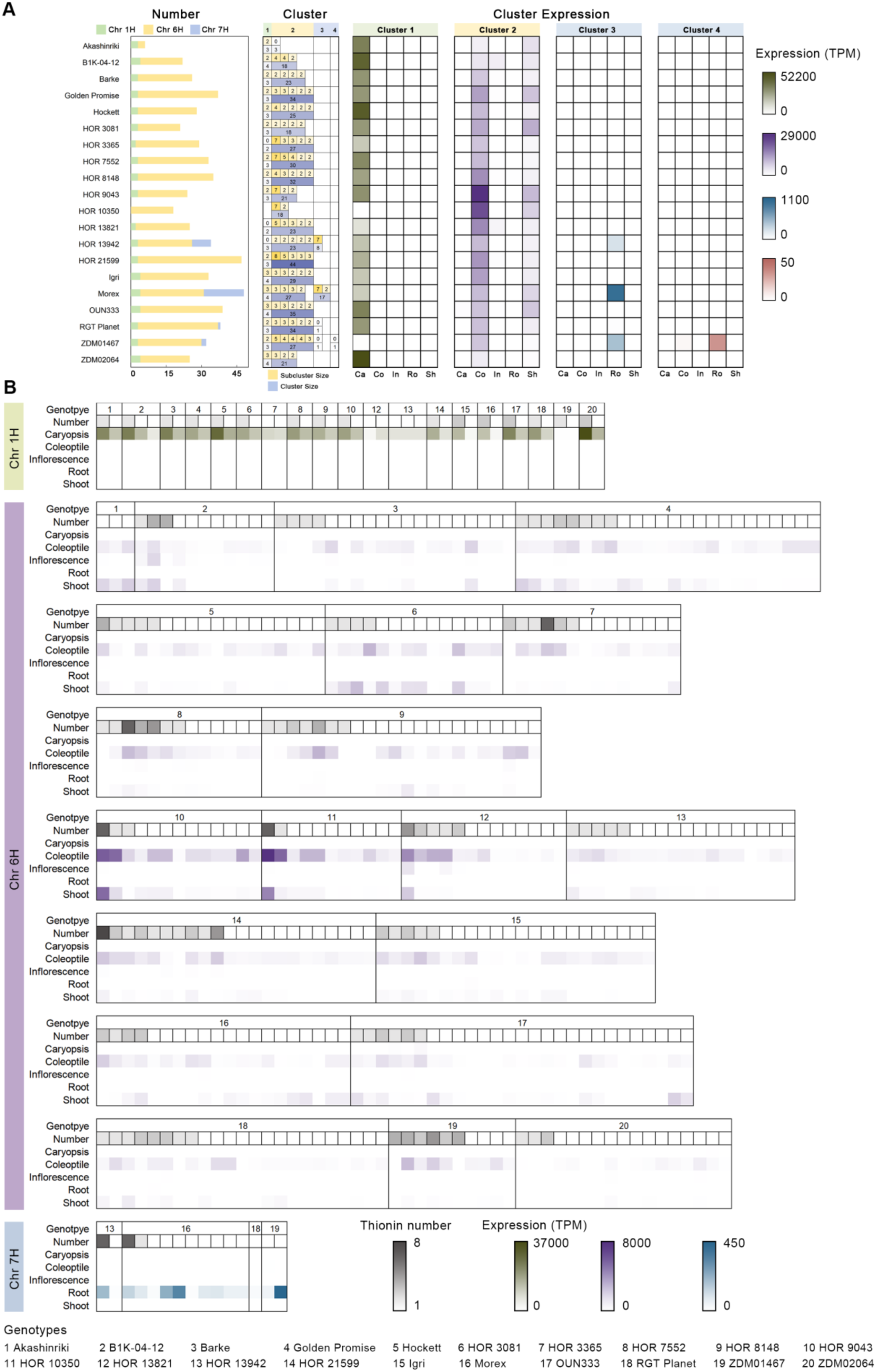
Analysis of thionin numbers and copy groups in the pan-transcriptome across 20 genotypes. (A) Thionin abundance, clustering structure, and tissue-specific expression patterns across 20 barley genotypes. The left panel shows the number of thionin transcripts detected in the pan-transcriptome for each genotype, grouped by chromosome. The middle panel summarizes the assignment of transcripts to conserved thionin clusters, with numbers indicating the size of each cluster and its corresponding subclusters. The right panel displays heatmaps of expression levels (transcripts per million, TPM) for clusters 1-4 across five tissues, including caryopsis (Ca), coleoptile (Co), inflorescence (In), root (Ro), and shoot (Sh). Color intensity reflects transcript abundance, with darker colors indicating higher expression levels. (B) Detailed expression profiles of thionins across genotypes and tissues, organized by chromosome. For each chromosome, individual genotypes are represented by numerical labels, which correspond to genotype names shown in the legend below. Within each genotype, single transcripts and subclusters are displayed as rows, and the number of transcripts within each subcluster is indicated by grayscale shading. Heatmaps illustrate the expression patterns of thionins across caryopsis, coleoptile, inflorescence, root, and shoot for different genotypes and chromosomes. Expression levels (TPM) are represented using color gradients. Detailed transcript and cluster information is provided in Table S14.

Pan-genome analysis identified 10 thionin clusters and 5 singleton sequences (Figure 4C, Table S10). Mapping these clusters to the pan-transcriptome showed that only clusters 1-4 were detected, while clusters 5-10 and all singleton sequences were absent (Table S13). These results indicate that thionin expression in the pan-transcriptome is mainly contributed by a small number of conserved and relatively large clusters. Distinct tissue-specific expression patterns were observed among thionin clusters located on different chromosomes (Figure 5A, B, Table S14). Cluster 1 on chr 1H exhibited strong caryopsis-specific expression across all genotypes, with expression levels reaching 10^4^-10^5^ transcripts per million (TPM) in caryopsis, while remaining close to background levels in other tissues. At the genotype level, ZDM02064 displayed the highest caryopsis expression, while ZDM01467 showed markedly reduced expression, indicating substantial inter-genotypic variation (Figure 5A). Although cluster 1 typically contained only two to four transcripts per genotype, its exceptionally high expression levels resulted from the cumulative contribution of these highly similar transcripts.

In contrast, cluster 2 on chr 6H displayed a markedly different expression profile. Expression was concentrated in coleoptile and shoot, while overall expression in caryopsis remained low (Figure 5A). This cluster contained the largest number of transcripts among all clusters, ranging from 18 to 44 per genotype, and exhibited high overall expression across tissues. Coleoptile expression was consistently high in all genotypes, with the highest level observed in HOR9043 (Figure 5A). The strongest shoot expression was detected in HOR3081. Thionins located on chr 7H, corresponding to clusters 3 and 4, showed the lowest overall expression levels but displayed a highly consistent root-specific expression pattern (Figure 5A). Among the detected genotypes, Morex exhibited the most pronounced root expression. Cluster 4 was detected exclusively in ZDM01467 and showed relatively low expression levels while remaining preferentially expressed in root. These results indicate that chr 7H thionins primarily function in belowground tissues.

Multiple transcripts within individual genotypes were found to be completely identical at the nucleotide sequence level. Expression levels could not be reliably distinguished at the individual transcript level under these conditions. Transcripts with 100% sequence identity were therefore grouped into subclusters (Table S13), and their expression values were combined. As shown in Figure 5A and 5B, multiple subclusters were identified in each genotype. HOR21599 contained the largest subcluster, comprising eight transcripts, and harbored the highest number of thionins on chr 6H. Its overall thionin expression level was not the highest among genotypes, indicating that expression intensity does not correlate with subcluster size or thionin copy number. Similar patterns were observed in other genotypes, suggesting that thionin expression is regulated by multiple factors rather than CNV alone.

Overall, thionins displayed pronounced chromosome-dependent and tissue-specific expression patterns at the pan-transcriptome level. Expression was strongest on chr 1H and highly concentrated in caryopsis. Intermediate expression levels were observed on chr 6H, with predominant expression in coleoptile and shoot. Expression on chr 7H was lowest and largely restricted to root. These results demonstrate a clear functional diversification of thionins across chromosomes and tissues in barley and provide a framework for further investigation of their roles in plant defense and development.

### Aphid infestation triggers genotype-dependent expression of chr 6H thionins in barley

Previous studies have demonstrated that thionin genes are strongly induced in barley following aphid infestation, with higher expression levels typically observed during interactions of aphids with poor-host plants (Delp et al., 2009; Mehrabi et al., 2014; Escudero-Martinez et al., 2017). Notably, two barley thionin genes located on chr 6H (AK252675.1/MLOC_46400.1 and AK359149/MLOC_34881.1) have been shown to contribute to quantitative resistance against aphids by reducing aphid feeding efficiency or performance (Escudero-Martinez et al., 2017). Given the high sequence similarity and putative functional redundancy among thionin family members located on the same chromosome, it is likely that multiple thionins on chr 6H collectively contribute to barley responses to aphid infestation. Pan-genome analysis revealed substantial variation in the number of thionin genes located on chr 6H among barley genotypes. Based on this variation, Akashinriki, HOR10350, Morex, and HOR21599, four representative genotypes were selected for further analysis, harboring 3, 19, 30, and 47 thionin genes on chr 6H, respectively.

Due to the high sequence similarity among thionin family members, we were unable to assess expression of individual genes. Therefore, genotype-specific consensus primers targeting all chr 6H thionins were designed to quantify their total expression levels (Table S15). Plants were infested with the *R.padi*, for which barley is a suitable host (host interaction), and with *M.persicae*, for which barley represents a poor host (poor-host interaction). Expression levels of chr 6H thionins were subsequently quantified by RT–qPCR (Figure 6, S1). Under untreated conditions, basal expression levels of chr 6H thionins differed markedly among the four genotypes (Figure 6, S1). HOR10350 consistently exhibited the highest basal expression, showing a clear separation from the other three genotypes, whereas Akashinriki displayed the lowest expression levels. Basal expression in HOR21599 was slightly lower than that in Morex. Interestingly, in the pan-transcriptome analysis of relative thionin expression across genotypes, all chr 6H thionins were assigned to cluster 2 . Comparison of cluster 2 expression levels among the four genotypes revealed the following ranking: HOR10350 > Morex > Akashinriki > HOR21599 (Figure 5A, Table S14). This overall expression pattern was broadly consistent with the relative expression trends obtained by RT-qPCR, although neither dataset showed a correlation between thionin expression levels and thionin gene copy number. Following aphid infestation, thionin expression changed in all four genotypes (Figure 6, S1). However, both the magnitude and direction of the response varied among genotypes and between aphid species. In Akashinriki, HOR10350, and HOR21599, thionin expression was consistently upregulated following infestation by both aphid species, with a stronger induction observed in response to *M. persicae* than to *R. padi*. In contrast, Morex displayed only minor changes in thionin expression following aphid infestation. Notably, in some biological replicates, thionin expression in Morex even showed a tendency toward downregulation after *M. persicae* infestation (Figure 6, S1). This unusual response pattern was consistent with previous observations (Escudero-Martinez et al., 2017). Comparison of thionin expression levels among genotypes further revealed pronounced genotype-dependent differences. HOR10350 consistently displayed the highest expression levels, followed by HOR21599 and Morex, whereas Akashinriki exhibited the lowest expression levels. Importantly, this expression ranking did not correspond to the relative differences in thionin gene copy number among genotypes, indicating that aphid-induced thionin expression is not solely determined by CNV (Figure 6, S1).

**FIGURE 6.**
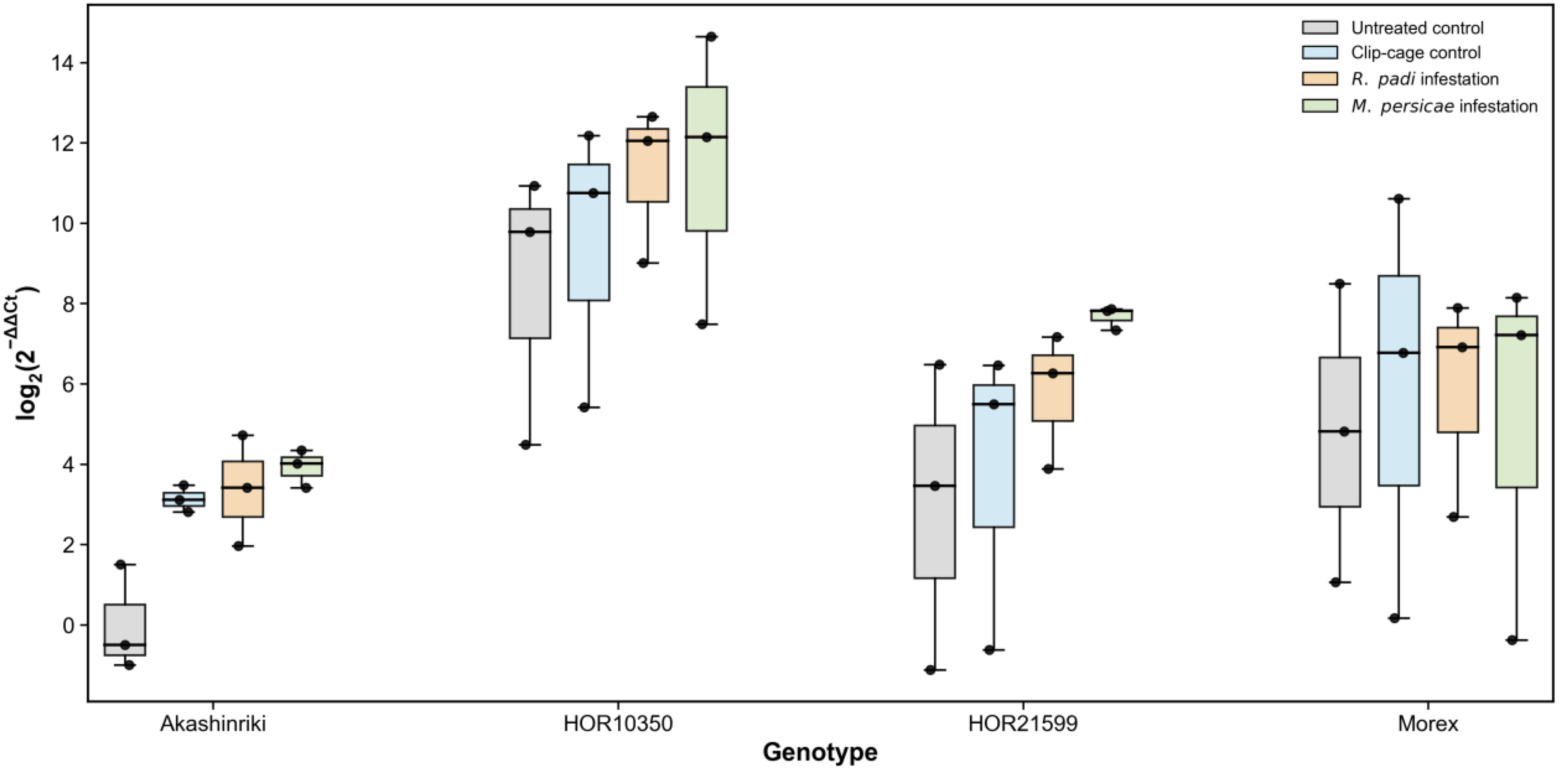
Relative expression levels of chr 6H thionins in four barley genotypes under different aphid infestation conditions. Relative expression levels of chr 6H thionins were determined by RT-qPCR in Akashinriki, HOR10350, HOR21599, and Morex under untreated control, clip-cage control, *R.padi* infestation, and *M.persicae* infestation conditions, *ACT* was used as the reference gene. Expression levels were calculated using the 2^-△△Ct^ method and are presented as log_2_-transformed values. Boxes indicate the interquartile range, horizontal lines within boxes represent the median, black dots indicate the mean, and whiskers represent the minimum and maximum values. Each point represents an independent biological replicate.

## DISCUSSION

### Evolutionary expansion of the barley thionin gene family

In this study, we systematically characterized the barley thionin gene family using the Morex V3 reference genome and identified 56 thionin genes, including eight genes with incomplete gene structures that nonetheless retain the conserved thionin domain. Phylogenetic analysis revealed a clear association between chromosomal location and evolutionary grouping: genes located on the same chromosome tend to form tightly clustered genomic regions with high sequence similarity, whereas genes located on different chromosomes belong to distinct phylogenetic clades. This pattern strongly suggests that tandem duplication has played a major role in the expansion of the thionin gene family in barley. Interestingly, thionins located on chr 5H and 7H were found to contain ten cysteine residues, exceeding the canonical number typically observed in classical thionin domains, suggesting that these genes may represent structurally divergent thionin variants.

CNV of the thionin gene family in barley has been recognized for several decades. Early Southern blot analyses estimated that the barley haploid genome contains at least 50-100 thionin gene copies (Bohlmann et al., 1988). Comparative hybridization analyses across different *Hordeum* species further revealed substantial variation in thionin copy numbers among species (Bunge et al., 1992). With the availability of genomic resources, multiple thionin-like genes were identified in the barley genome, many of which occur in clustered arrangements along chromosomes (Mascher et al., 2017).

Notably, the extensive and relatively homogeneous tandem expansion of thionin genes observed in barley appears to be uncommon in other plant species. For instance, only four thionin genes have been identified in Arabidopsis, whereas the thionins reported in rice exhibit greater sequence divergence rather than the large-scale tandem duplication pattern observed in barley (Epple et al., 1997; Boonpa et al., 2019). In wheat, thionins form a multigene family distributed across the homoeologous chromosomes of the A, B, and D genomes, although their total copy number has not yet been systematically determined (Castagnaro et al., 1992).

### CNV generates structural diversity in barley thionins

CNV represents an important source of genetic diversity and plays a critical role in trait evolution, environmental adaptation, and genome evolution. It refers to variations in the copy number of specific DNA segments within the genome, resulting in the gain or loss of genes or sequences in particular genomic regions (Żmieńko *e*t al., 2014; Lye et al., 2019; Silaiyiman et al., 2025).

In plants, CNVs are widespread and frequently enriched in gene families associated with defense and stress responses. Genome-wide studies in maize, Arabidopsis, rice, and wheat have demonstrated that CNVs can arise over relatively short evolutionary timescales and influence gene expression as well as phenotypic diversity (Springer et al., 2009; DeBolt, 2010; Yu et al., 2013; Jiao et al., 2020; Żmieńko et al., 2020). In barley, both reference genome and pan-genome analyses have revealed extensive structural variation, particularly within gene clusters associated with defense-related functions (Mascher et al., 2017; Jayakodi et al., 2024). More recent pan-genome analyses have further identified the thionin locus as a prominent CNV hotspot, with copy numbers ranging from three to more than seventy across barley genotypes (Jayakodi et al., 2024). Notably, the 20 barley genotypes analyzed in this study were included in previously published barley pan-genome and pan-transcriptome resources designed to capture representative global barley diversity, including cultivars, landraces, and wild accessions originating from diverse geographic regions and contrasting growth habits (Jayakodi et al., 2024; Guo et al., 2025). These representative resources therefore provide a robust framework for investigating thionin structural and transcriptional diversification across barley germplasm. Using these datasets, we found that many thionin genes are conserved across genotypes and often share extremely high sequence similarity. The chromosomal distribution pattern was highly consistent across genotypes: thionins located on chr 1H were relatively conserved, typically comprising only two to four members per genotype, whereas chr 6H and 7H were enriched for thionin genes and exhibited pronounced CNV. Clustering analysis based on CDS identity identified ten clusters and five singleton sequences, with all chr 1H thionins belonging to a single cluster, while those on chr 6H and 7H were distributed across multiple clusters. Integration of pan-transcriptome data further showed that transcriptional evidence was largely restricted to several major clusters, whereas other minor clusters or singleton sequences showed no detectable expression under the sampled conditions. This pattern suggests that a limited set of core thionin clusters may account for most functional expression, while other members may represent a genetic reservoir for future functional diversification.

### Chromosome-specific expression divergence of thionin genes

CNV has been widely reported to influence gene function and phenotypic variation in plants (McHale et al., 2012). In several cases, CNVs have been directly linked to agronomic traits and environmental adaptation. For example, CNVs at the GL7 locus contribute to grain size diversity in rice (Wang et al., 2015), while variation in the copy number of Ppd-B1 modulates flowering time in wheat (Würschum et al., 2015). Similarly, copy number variation in the maize transporter gene MATE1 affects aluminum tolerance (Maron et al., 2013). In barley, CNVs are also widespread across the genome (Bretani et al., 2020) and have been associated with adaptive traits and stress tolerance. For instance, increased copy number of the efflux transporter BOR1 confers tolerance to boron toxicity (Sutton et al., 2007), and variation within the HvCBF4-HvCBF2 gene cluster has been shown to significantly affect frost tolerance (Francia et al., 2016). Despite these well-characterized examples, the functional consequences of the extensive gene CNV present in plant genomes remain incompletely understood.

In this study, we characterized CNV patterns of the thionin gene family across 20 barley genotypes and integrated sequence- and expression-based analyses to explore their potential functional diversification. Clear chromosome-associated differences were observed in both copy numbers and expression patterns. Thionins located on chr 1H exhibited the lowest copy numbers and the smallest cluster sizes and were predominantly expressed in caryopsis tissues with consistently high transcript levels under non-stressed conditions. This pattern suggests relatively conserved functions and a potential role in seed development. In contrast, thionins on chr 6H showed substantially higher copy numbers and extensive cluster expansion, indicative of recurrent local duplication events. These genes were mainly expressed in coleoptile and shoot tissues and, based on previous studies, are likely associated with antimicrobial defense and resistance to insect pests (Höng et al., 2021; Li et al., 2021). Thionins on chr 7H were also numerous at the genomic level, however, only a limited number of sequences and genotypes showed detectable expression in the pan-transcriptome, primarily in root tissues. For most 7H thionins, no transcriptional evidence was observed under the sampled conditions, suggesting that their expression may be restricted to specific tissues not represented in the current dataset or may require induction by environmental or biotic stress signals. Collectively, these results indicate that CNV within the barley thionin family is associated with pronounced expression divergence across chromosomes, supporting distinct functional roles in different tissues and developmental contexts.

Using the available pan-transcriptome data, we did not detect a direct relationship between thionin copy number, cluster size, or subcluster size and transcript abundance. This apparent decoupling may reflect the role of thionins as components of the plant immune system, whose expression is often tightly regulated and primarily activated under stress conditions. Previous functional studies of barley thionins have also been limited by the high sequence similarity among family members, which complicates the isolation and characterization of individual genes. In this context, the Akashinriki genotype represents a particularly valuable resource, as it contains only three thionin genes on chr 6H, each with a distinct sequence. This simplified genetic architecture provides a useful entry point for dissecting the specific functions of 6H thionins. Similarly, chr 1H contains only two to four thionin genes across genotypes, including one unique gene and a small duplicated group, which may further facilitate functional analysis. Notably, the genotype HOR10350 lacks thionin genes on chr 1H altogether. Whether this absence affects specific aspects of plant development or stress responsiveness remains an intriguing question and warrants further investigation.

### Thionins as components of plant stress and defense signaling

To explore the potential functions of the thionin gene family in barley, we analyzed *cis*-elements in the promoter regions of 56 thionin genes in the barley reference genome and examined their expression patterns across 20 genotypes using pan-transcriptomic data. These analyses revealed pronounced expression specificity among thionins located on different chromosomes. Notably, thionins located on chr 6H were predominantly expressed in coleoptile and shoot tissues, consistent with their historical designation as leaf thionins (Bohlmann et al., 1988).

Promoter *cis*-element analysis of thionin genes in Morex V3 revealed a high abundance of regulatory elements associated with plant defense and stress-related hormone signaling. In particular, *cis*-elements linked to ABA, SA, and MeJA pathways were frequently identified. These hormone-signaling pathways play central roles in plant immunity and stress responses (Bari et al., 2009; Pieterse et al., 2012). Consistent with this, previous studies have shown that barley thionin expression is strongly induced by fungal and bacterial pathogens (Stevens et al., 1996). For example, a pronounced accumulation of thionins was observed following powdery mildew infection (Bohlmann et al., 1988), and heterologous expression of barley thionin genes in *Nicotiana tabacum* L. conferred enhanced resistance to pathogens (Carmona et al., 1993). Together, these findings support a conserved role for barley thionins in plant defense mechanisms.

In addition to pathogen resistance, hormone-mediated signaling pathways are also central to plant defense against phloem-feeding insects. Jasmonic acid and salicylic acid signaling have been shown to regulate plant responses to aphid infestation (Zhu-Salzman et al., 2004; Walling, 2008; Morkunas et al., 2011; Avila et al., 2012). In barley, analysis of the partially resistant wild barley Hsp5 revealed that resistance traits were primarily associated with mesophyll and phloem tissues and were accompanied by elevated ABA- and JA-related signaling following aphid infestation (Leybourne et al., 2019). Moreover, the involvement of ABA signaling in aphid resistance has been well established (Thaler et al., 2004). Consistently, transgenic barley plants expressing the aphid effector Rp1, which enhances host susceptibility, exhibited significant downregulation of genes associated with SA and JA signaling pathways (Escudero-Martinez et al., 2020). The enrichment of ABA-, SA-, and JA-related *cis*-elements identified in this study aligns with these observations and suggests that thionins may contribute to barley resistance against aphids through regulation by key hormone signaling networks. Interestingly, a substantial number of light-responsive *cis*-elements were also detected in the promoter regions of barley thionin genes, indicating potential regulation of thionin expression by light. This finding is consistent with earlier work by Reimann-Philipp et al (1989), who reported a rapid decrease in leaf thionin-specific mRNA levels in barley seedlings following sudden exposure to light. This response was proposed to involve interactions between multiple light receptors. However, how light-mediated repression of thionin expression is integrated with stress- or defense-induced activation remains unclear and warrants further investigation.

### Induction of thionins during barley-aphid interactions

Previous studies have demonstrated that aphid infestation induces the expression of thionin genes in barley (Escudero-Martinez et al., 2017; Leybourne et al., 2019; Escudero-Martinez et al., 2020). However, functional characterization of individual thionin genes has long been hampered by limited knowledge of thionin gene number, copy composition, and precise sequence boundaries. The high sequence similarity among thionin family members has further complicated the reliable discrimination and analysis of specific genes. By integrating pan-genome and pan-transcriptome resources, the present study systematically resolved thionin numbers across diverse barley genotypes and characterized their basal expression profiles under natural growth conditions. This comprehensive framework provided a robust foundation for examining thionin responses to aphid infestation.

Given the pronounced variation in thionin number on chr 6H among genotypes, 4 representative genotypes spanning a clear gradient of chr 6H thionin number were selected for aphid infestation assays. Because of the exceptionally high sequence similarity among thionins localized on chr 6H, gene-specific primers could not reliably distinguish individual members, even with careful design. Consequently, chr 6H thionins were analyzed as a single gene set within each genotype, and genotype-specific consensus primers were designed. To ensure specificity and robustness, primers were derived exclusively from conserved regions shared among full-length thionin sequences. As all chr 6H thionins belonged to cluster 2, the primers effectively targeted conserved regions of cluster 2 thionins across genotypes.

Expression analyses following aphid infestation confirmed that thionins were strongly induced by aphid feeding, with consistently higher induction observed in poor-host interactions. This finding suggests that thionins participate in barley defense responses to aphids and offers a potential molecular explanation for the enhanced resistance typically observed in poor-host genotypes. Comparative analyses revealed genotype-specific thionin expression patterns: HOR10350 maintained high basal expression, HOR21599 showed the strongest induction following aphid infestation, whereas Akashinriki exhibited consistently low expression, in line with its lower thionin copy number on chr 6H. These observations suggest that the level of thionin gene induction in response to aphid infestation may be positively associated with thionin number, even though basal expression levels do not correlate with gene family size.

Notably, Morex exhibited a distinct expression pattern compared with the other genotypes, showing little induction or even a reduction in thionin expression following aphid infestation. Similar atypical responses have been reported previously (Escudero-Martinez et al., 2017), and the underlying causes remain unclear. These patterns may reflect genotype-specific regulatory mechanisms, differences in signaling network integration, or broader genomic background effects, and would require further investigation.

Taken together, under natural growth conditions, thionin expression levels did not show a clear relationship with gene number. In contrast, under aphid infestation, genotype-dependent changes in thionin expression tended to increase with number. These findings indicate that thionins contribute to genotype-specific aphid defense responses in barley. Nevertheless, aphid resistance is a highly complex trait involving multiple defense components and signaling pathways. How thionins are integrated into this broader defense network, and how they interact with other resistance factors, remains to be explored.

## AUTHOR CONTRIBUTIONS

**Yao Fu**: Experimental design; data curation; formal analysis; investigation; methodology; visualization; writing—original draft; writing—review and editing. **Joanne Russell:** Conceived and directed the project; writing—review and editing. **Miriam Schreiber:** Methodology; analytical supervision; strategic guidance; writing—review and editing. **Jorunn I. B. Bos:** Funding acquisition; conceived and directed the project; experimental design; writing—review and editing.

## ACKNOWLEDGMENTS

The authors thank Research Computing at the James Hutton Institute for providing computational resources and technical support for the “UK’s Crop Diversity Bioinformatics HPC” (BBSRC grants BB/S019669/1 and BB/X019683/1). JR and MS acknowledge funding from the Scottish Government’s Rural and Environmental Science and Analytical Services (RESAS) strategic research programme under BARGAIN (JHI-B1-2). YF acknowledges support from the China Scholarship Council (CSC).

## DATA AVAILABILITY STATEMENT

All datasets used in this manuscript are publicly available. The code used for data processing and analysis is available at https://zenodo.org/records/20433719.

## SUPPLEMENTAL MATERIAL

**Table S1.** Coding and protein sequences of the barley thionin gene family identified in the Morex V3 genome

**Table S2.** Physicochemical characteristics and signal peptide predictions of barley thionin proteins in the Morex V3 genome

**Table S3.** Conserved domain annotations of barley thionin proteins identified in the Morex V3 genome

**Table S4.** Chromosomal locations and distribution of barley thionin genes in the Morex V3 genome

**Table S5.** Pairwise nucleotide and amino acid sequence identity matrices of barley thionins in the Morex V3 genome

**Table S6.** *Cis*-regulatory element composition of promoter regions of barley thionin genes in the Morex V3 genome

**Table S7.** Functional classification and annotation of identified *cis*-regulatory elements

**Table S8.** Coding and protein sequences of thionins across 20 barley genotypes in the pan-genome

**Table S9.** Pairwise sequence identity analysis of thionins across 20 barley genotypes in the barley pan-genome

**Table S10.** Clustering and subclustering of thionin sequences across 20 barley genotypes in the barley pan-genome

**Table S11.** Genetic *p*-distances among thionin clusters and singleton sequences across 20 barley genotypes

**Table S12.** Transcript and protein sequences of thionins identified in the barley pan-transcriptome

**Table S13.** Clustering and subclustering of thionin transcripts across 20 barley genotypes in the barley pan-transcriptome

**Table S14.** Expression profiles of barley thionin genes across tissues and experimental conditions

**Table S15.** Primers used for RT-qPCR validation of thionin gene expression **Figure S1.** Relative expression levels of chromosome 6H thionin genes in four barley genotypes under different aphid infestation treatments, using *UBC* as the reference gene

## Abbreviations

*ACT*: actin
ABA: abscisic acid
CDS: coding sequence
CNV: copy number variation
HMM: hidden Markov model
HSP5: *Hordeum spontaneum* 5
JA: jasmonic acid
MeJA: methyl jasmonate
SA: salicylic acid
TPM: transcripts per million
*UBC*: ubiquitin-conjugating enzyme.

